# Tailoring and optimizing fatty acid production by oleaginous yeasts through the systematic exploration of their physiological fitness

**DOI:** 10.1101/2022.06.20.496586

**Authors:** Zeynep Efsun Duman-Özdamar, Vitor A.P. Martins dos Santos, Jeroen Hugenholtz, Maria Suarez-Diez

**Affiliations:** Bioprocess Engineering, Wageningen University & Research, 6708 PB, Wageningen, the Netherlands; Laboratory of Systems and Synthetic Biology, Wageningen University & Research, 6708 WE, Wageningen, the Netherlands; LifeGlimmer GmbH, Berlin, 12163, Germany; Wageningen Food & Biobased Research, Wageningen University & Research, 6708 WE, Wageningen, The Netherlands

**Keywords:** Oleaginous yeasts, microbial oil, Response Surface Methodology, carbon to nitrogen ratio, lipid accumulation

## Abstract

**Background:** The use of palm oil for our current needs is unsustainable. Replacing palm oil with oils produced by microbes through the conversion of sustainable feedstocks is a promising alternative. However, there are major technical challenges that must be overcome to enable this transition. Foremost among these challenges is the stark increase in lipid accumulation and production of higher content of specific fatty acids. Therefore, there is a need for more in-depth knowledge and systematic exploration of the oil productivity of the oleaginous yeasts. In this study, we cultivated *Cutaneotrichosporon oleaginosus* and *Yarrowia lipolytica* at various C/N ratios and temperatures in a defined medium with glycerol as carbon source and urea as nitrogen source. We ascertained the synergistic effect between various C/N ratios of a defined medium at different temperatures with Response Surface Methodology (RSM) and explored the variation in fatty acid composition through Principal Component Analysis.

**Results:** By applying RSM, we determined a temperature of 30 °C and a C/N ratio of 175 g/g to enable maximal oil production by *C. oleaginosus* and a temperature of 21 °C and a C/N ratio of 140 g/g for *Y. lipolytica*. We increased production by 71 % and 66 % respectively for each yeast compared to the average lipid accumulation in all tested conditions. Modulating temperature enabled us to steer the fatty acid compositions. Accordingly, switching from higher temperature to lower cultivation temperature shifted the production of oils from more saturated to unsaturated by 14 % in *C. oleaginosus* and 31 % in *Y. lipolytica*. Higher cultivation temperatures resulted in production of even longer saturated fatty acids, 3 % in *C. oleaginosus* and 1.5 % in *Y. lipolytica*.

**Conclusions:** In this study, we provided the optimum C/N ratio and temperature for *C. oleaginosus* and *Y. lipolytica* by RSM. Additionally, we demonstrated that lipid accumulation of both oleaginous yeasts was significantly affected by the C/N ratio and temperature. Furthermore, we systematically analyzed the variation in fatty acids composition and proved that changing the C/N ratio and temperature steer the composition. We have further established these oleaginous yeasts as platforms for production of tailored fatty acids.

## Introduction

The use of plant-derived oils, especially palm oil, is increasing at an alarming rate. This is happening in part as a replacement for fossil foils, but mostly as they are cheap sources of many useful components. The oils and fatty acids derived from palm trees are used in food, feed, chemical, personal care, and cosmetic products for health benefits, sensorial reasons (texture, flavor), to extend shelf-life, and as surfactants or emulsifiers [1–3]. As a result, palm tree groves are rapidly replacing the original tropical forests, and other original and traditional vegetation in many Asian, South American, and African countries. This replacement is not only threatening the local ecosystem but is also having a major effect on the local livelihoods, as it causes deforestation and contributes to climate change [4,5]. Despite some responsible actions that have been taken, among them fighting against deforestation driven by RSPO (Roundtable on Sustainable Palm Oil), the use of palm oil remains controversial [6]. To that end, developing a sustainable alternative to fatty acids and oils is urgent and of utmost interest.

Oil-producing yeasts, referred to as oleaginous yeasts, have strong potential as sustainable alternatives for lipid production in various industrial applications [7]. *Yarrowia lipolytica* and *Cutaneotrichosporon oleaginosus* also known as *Apiotrichum curvatum, Cryptococcus curvatus, Trichosporon cutaneum, Trichosporon oleaginosus*, and *Cutaneotrichosporon curvatum* are reported among the top five most well-known oil-producing yeasts. *C. oleaginosus* and *Y. lipolytica* can accumulate oils up to 70% and 40% of their biomass respectively [8–10]. Lipid accumulation in oleaginous yeasts is induced by limiting specific nutrients such as nitrogen, phosphate, and sulphur. Nitrogen limitation or, in other words, a high C/N ratio in the growth medium, has been observed to be the most effective lipid induction strategy [9]. As reported by Ykema et al., after passing the critical C/N of 11 g/g, the oleaginous yeast starts to accumulate oils by re-routing the excess carbon to be stored as lipids [11,12]. Under nitrogen limiting conditions, the produced fatty acid composition has been reported to be 25% palmitic acid (C16:0), 10% stearic acid (C18:0), 57% oleic acid (C18:1), and 7% linoleic acid (C18:2) by *C. oleaginosus* [13] and 15% C16:0, 13% C18:0, 51% C18:1, and 21% C18:2 by *Y. lipolytica* [14]. This composition is comparable to that of palm oil. Furthermore, these yeasts can use a broad range of carbon sources, such as glucose, xylose, glycerol, sucrose, and lactose. They can also use more complex and inexpensive side streams such as crude glycerol from bioethanol production or whey permeate as a feedstock, which is significant to reduce raw materials cost [15–17]. Moreover, *Y. lipolytica* is non-pathogenic and regarded as food-grade yeast, thus its oil can be used for food-related applications [18,19]. Due to these advantages, oleaginous yeasts are flagged as attractive microbial-cell factories to sustain a bio-based circular economy for industrial implementation. However, from an economic point of view, the lipid production process of oleaginous yeasts still requires substantial optimization for industrial purposes. Koutinas et al. [20] reported that implementation of microbial oil in industrial applications is strongly dependent on the final microbial oil concentrations and lipid productivity [19]. For *C. oleaginosus*, it has been calculated that the process is only economically feasible if lipid accumulation reaches approximately 85 % (w/w).

Identifying and designing optimal production conditions is a challenging step in developing bioconversion systems since these cultivation conditions play a crucial role in productivity [21]. For at least two decades, efforts have been made to design the optimum growth medium and fermentation conditions to boost the lipid accumulation as well as to sustain the growth of oleaginous yeasts [22–24]. One of the most efficient strategies to systematically identify optimum production conditions is through the Response Surface Methodology (RSM). This method decreases experimental time and laborious work compared to the one-factor-at-a-time (OFAT). Whereas OFAT only allows changing one of the considered factors in each of the experiments, RSM provides a design on which multiple parameters are changed at each experimental run. Moreover, RSM aims to predict the observed response, by reliably estimating the experimental variability [25]. For instance, Awad et al. assessed the effect of various carbon and nitrogen sources on the physiology of *C. oleaginous* via RSM [26]. In another example, Cui et al. focused on the effect of temperature and pH on lipid content and the growth on crude glycerol [27]. Additionally, Canonico et al. reported optimum C/N ratio and time to maximize lipid production of *Y. lipolytica* [17]. However, no studies have focussed on the synergistic effect between the C/N ratio of a defined medium and cultivation parameters.

In addition to the optimized lipid productivity of oleaginous yeasts, tailoring the composition of produced lipids is important for increasing economic competitiveness. For instance, the longer fatty acids, such as oleic acid, lauric acid, and palmitic acid are heavily used in home and personal care products due to their cleaning/surfacting activities. Polyunsaturated fatty acids (PUFAs) such as linolenic acid supply various health properties. Ochsenreither et al. [19] reported that temperature and the composition of the medium lead to variation in fatty acid composition. Therefore, analyzing the fatty acid compositions under different cultivation conditions will provide us valuable information that will enable us to devise strategies to tailor fatty acid compositions.

Thus, in this study, we aimed to further develop *C. oleaginosus* and *Y. lipolytica* as microbial cell factories for the improved production of lipids and lipids with higher content of specific fatty acids. The first set of experiments was designed to assess the variance between the fatty acid composition produced by *C. oleaginosus* and *Y. lipolytica* at various C/N ratios and temperatures. We extensively studied the variance in fatty acids composition via PCA. Additionally, we performed another set of experiments to broaden the current experimental region for RSM and determined the optimized C/N ratio and temperature for *C. oleaginosus* and *Y. lipolytica*.

## Materials and Methods

### Yeast Strains and Pre-culture Preparation

*Cutaneotrichosporon oleaginosus* ATCC 20509 and *Yarrowia lipolytica* DSM 1345 were maintained on Yeast extract Peptone Dextrose (YPD) agar plates containing 10 g/L yeast extract, 20 g/L peptone, 20 g/L glucose, 20 g/L agar. The maintained cultures were stored at 4 °C for up to a week. The inoculum was prepared by transferring a single colony of the oleaginous yeasts into 10 mL YPD broth (10 g/L yeast extract, 20 g/L peptone, 20 g/L glucose, 20 g/L) in 50 mL tubes and incubated at 30 °C, 250 rpm for 18 h in a shaking incubator.

### Cultivation Conditions

*C. oleaginosus* and *Y. lipolytica* were grown in defined media consisting of glycerol as a carbon source and urea as a nitrogen source. The medium was adapted from Meester et al. [28] with modifications. In addition to the variation in carbon to nitrogen ration, the medium contained 2.7 g/L KH_2_PO_4_, 1.79 g/L NaH_2_PO_4_.7H_2_O, 0.2 g/L MgSO_4_·7H_2_O, 0.2 g/L MgSO_4_.7H_2_O, 0.1 g/L EDTA with the pH 5.5 as well as the trace elements: 40 mg/L CaCl_2_·2H_2_O, 5.5 mg/L FeSO_4_.7H_2_O, 5.2 mg/L citric acid.H_2_O, 1 mg/L ZnSO_4_·7H_2_O, 0.76 mg/L MnSO_4_.H_2_O, 10 μL/L H_2_SO_4_ (36N).

### Experimental Design

The C/N ratio varied from 30:1 (g/g) to 300:1 (g/g) by mixing 4 – 40 g carbon/L with 0.13 g nitrogen/L for both oleaginous yeasts. After that, prepared cultures were incubated at different temperatures (15 °C, 25 °C, 30 °C, and 35 °C), 250 rpm for 96 or 144 hours. For the experimental design, the C/N ratio (X_1_) and temperature (X_2_) were selected as variables. Low levels and high levels for C/N ratio and temperature are C/N 30, C/N 120, and 15 °C, 35 °C. The coded values of independent variables were calculated by considering the high and low levels together with the real values:

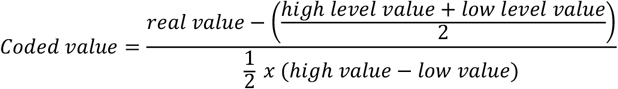

and are presented in Table 1.

**Table 1.** Levels of two independent variables employed in RSM in terms of real and coded values for *C. oleaginosus* and *Y. lipolytica*.

| Levels and coded values | Real values |  |
| --- | --- | --- |
|  | C/N ratio (g/g)<br>X<sub>1</sub> | Temperature (°C)<br>X<sub>2</sub> |
| -1.0 | 30 | 15 |
| -0.3 | 60 | - |
| 0.0 | 75 | 25 |
| 0.5 | - | 30 |
| 1.0 | 120 | 35 |
| 3.7 | 240 | - |
| 5.0 | 300 | - |

### Determination of Biomass

The growth of oleaginous yeasts was monitored every 24 h by measuring the absorbance at 600 nm (OD_600_). A calibration curve was plotted with the absorbance versus the dry cell weight for *C. oleaginosus* and *Y. lipolytica* (Figure S1). The dry cell was obtained as follows: (1) 5 mL of culture was centrifuged at 3200 g for 15 mins, (2) the cells were washed twice with 10 mL of deionized water (3) and freeze-dried.

### Identification of lipids and fatty acid composition

The total fatty acids and the fatty acid composition were determined quantitatively. The samples were prepared by mixing 20-25 mg freeze-dried yeast cells with 2 mL of 15% H_2_SO_4_ in methanol and 2 mL of chloroform containing methyl pentanoate as an internal standard. The samples were incubated for 4 h at 85-95 °C and cooled on ice for 5 min, 1 mL of distilled water was added. Following the phase separation by centrifugation at 2200 g for 5 min, the organic phase was collected from the bottom of the tube and dried with NaSO_4_. Subsequently, The fatty acid methyl esters (FAME) were analyzed with a gas chromatograph (Brand, City, Country) equipped with a Zebron ZB-FAME column (30 m x 0.25 mm x 0.20 μm; Phenomenex, Torrance, CA, The US). The yeast’s oil content was calculated from the internal standard.

### Computational Analysis

All computational analysis was performed with R version 4.0.2 [29].

*Principal Component Analysis* (*PCA*) was carried out on the fatty acid profiles by the statistics function prcomp within R [30]. The correlation biplots of the principal component scores and the loading vectors were plotted through R ggplot2 package [31].

*Response Surface Methodology* (*RSM*) was performed by the rsm package, and the contour plots were generated with the R pers or cont functions [32]. The relationship between the responses and factors was expressed by the second-order polynomial equation:

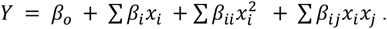

*Y* represents the predicted response, *βo* is the interception coefficient, *βi* is the linear coefficient, *β_ii_* is the quadratic coefficient and *β_ij_* is the interaction coefficient, X_i_ is the independent variable, X_i_^2^ is the squared effect, and X_i_X_j_ is the interaction effect. The quality of the regression equations was assessed according to the coefficient of determination (R^2^) and lack of fit F-test. Statistical analysis of the model was performed using Analysis of Variance (ANOVA) and p < 0.05 was considered significant. Optimal levels of the C/N ratio and temperature were given as stationary points via RSM.

## Results

In total, 10 experiments were conducted for *C. oleaginosus* and 9 experiments for *Y. lipolytica*.The biomass concentration of *C. oleaginosus* and *Y. lipolytica* varied from 1.24 g/L to 5.54 g/L and 2.34 to 4.52 g/L, lipid content ranged from 3.62 % to 47.41 % and 3.35 % and 18.51 % for all tested conditions (Tables 2 and 3). While the highest lipid content was obtained at C/N 120 at 30 °C for *C. oleaginous*, it reached the maximum point at C/N 140 at 25 °C in *Y. lipolytica*.

**Table 2.**
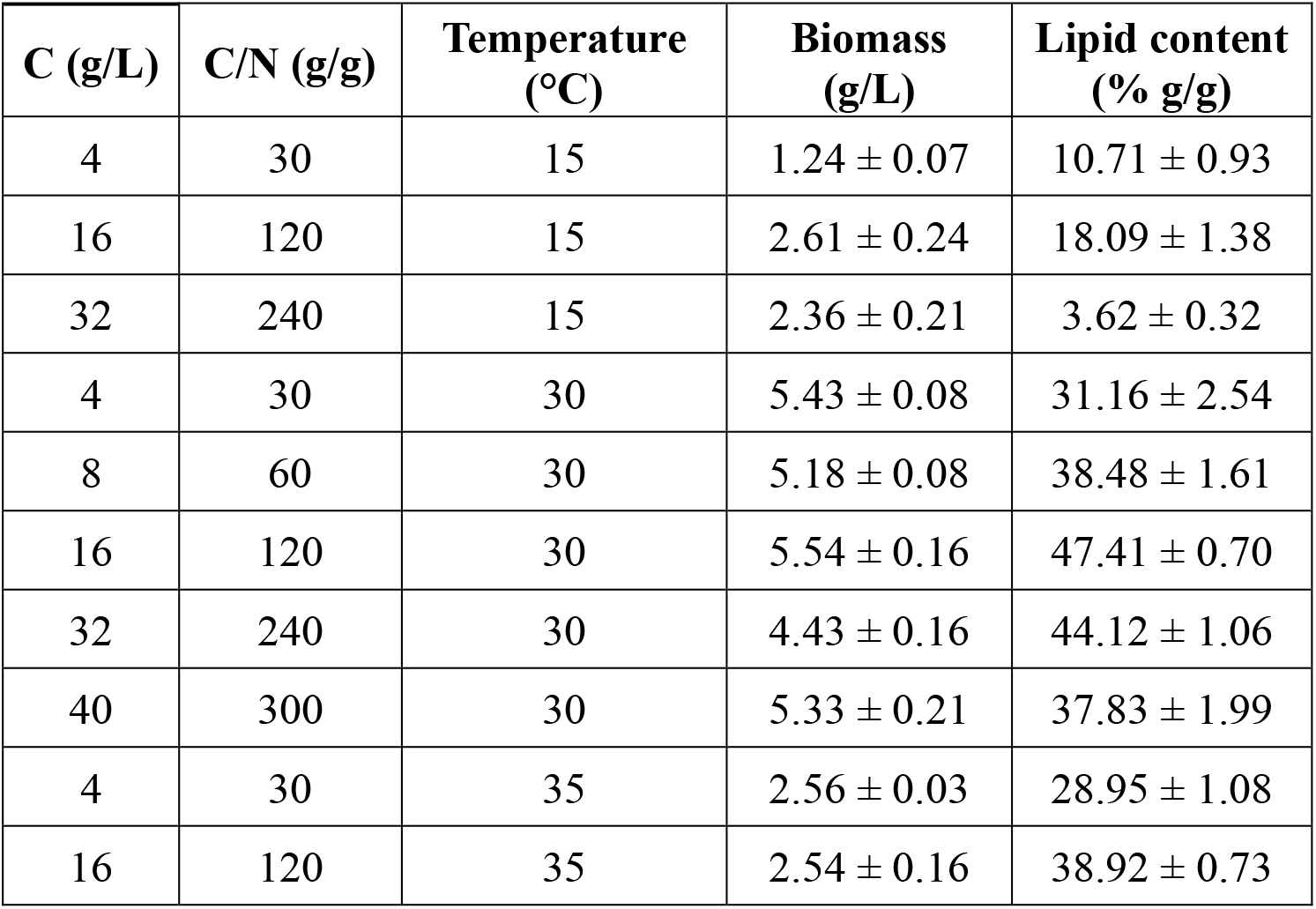
Biomass density and lipid content of *C. oleaginosus* at different C/N ratios and temperatures.

| C (g/L) | C/N (g/g) | Temperature (°C) | Biomass (g/L) | Lipid content (% g/g) |
| --- | --- | --- | --- | --- |
| 4 | 30 | 15 | 1.24 ± 0.07 | 10.71 ± 0.93 |
| 16 | 120 | 15 | 2.61 ± 0.24 | 18.09 ± 1.38 |
| 32 | 240 | 15 | 2.36 ± 0.21 | 3.62 ± 0.32 |
| 4 | 30 | 30 | 5.43 ± 0.08 | 31.16 ± 2.54 |
| 8 | 60 | 30 | 5.18 ± 0.08 | 38.48 ± 1.61 |
| 16 | 120 | 30 | 5.54 ± 0.16 | 47.41 ± 0.70 |
| 32 | 240 | 30 | 4.43 ± 0.16 | 44.12 ± 1.06 |
| 40 | 300 | 30 | 5.33 ± 0.21 | 37.83 ± 1.99 |
| 4 | 30 | 35 | 2.56 ± 0.03 | 28.95 ± 1.08 |
| 16 | 120 | 35 | 2.54 ± 0.16 | 38.92 ± 0.73 |

**Table 3.** Biomass density and lipid content of *Y. lipolytica* at different C/N ratios and temperatures.

| C (g/L) | C/N (g/g) | Temperature (°C) | Biomass (g/L) | Lipid content (% g/g) |
| --- | --- | --- | --- | --- |
| 4 | 30 | 15 | 2.62 ± 0.08 | 7.91 ± 0.49 |
| 16 | 120 | 15 | 4.40 ± 0.08 | 15.91 ± 0.07 |
| 4 | 30 | 30 | 2.45 ± 0.16 | 4.00 ± 0.24 |
| 16 | 120 | 30 | 4.52 ± 0.14 | 14.26 ± 1.09 |
| 4 | 30 | 35 | 2.34 ± 0.11 | 3.35 ± 0.33 |
| 16 | 120 | 35 | 3.12 ± 0.13 | 9.25 ± 0.06 |
| 9.75 | 75 | 25 | 3.01 ± 0.13 | 15.29 ± 0.41 |
| 9.75 | 75 | 25 | 3.06 ± 0.09 | 15.01 ± 0.17 |
| 18.2 | 140 | 25 | 4.18 ± 0.14 | 18.51 ± 0.96 |

Incubating *C. oleaginosus* at 15 °C slightly decreased the biomass and lipid content compared to the other tested temperatures. On the other hand, decreasing the C/N ratio reduced the biomass concentration of *Y. lipolytica*.

### Development of Regression Models and ANOVA

C/N ratio and temperature as coded variables, and lipid content and biomass as responses were analyzed with RSM. The relation was fitted by second-order polynomial equations to obtain the regression equation models (Table 4). These models represent the empirical relationships between the biomass density, lipid content of cells, and the variables (C/N ratio (*X*_1_) and temperature (*X*_2_)) in coded units. These regression models suggest that both linear and quadratic effects of C/N ratio and temperature significantly affected the lipid content of *C. oleaginosus* and *Y. lipolytica* (Table 4). However, interaction of C/N ratio and temperature has no significant effect on any considered responses of *Y. lipolytica*. When the growth of *C. oleaginosus* is not significantly affected by the C/N ratio, the combined effect of the C/N ratio and temperature has a significant effect. On the other hand, only the linear effect of investigated factors significantly affected the growth of *Y. lipolytica*.

**Table 4.** Regression equations, statistics of regression equations for lipid content and biomass of *C. oleaginosus* and *Y. lipolytica*.

| *C. oleaginosus* |  |  |  | *Y. lipolytica* |  |  |  |
| --- | --- | --- | --- | --- | --- | --- | --- |
| Dependent variable: Lipid content % (w/w) |  |  |  |  |  |  |  |
| $Y(\text{Lipid content})$<br>$= 39.26 + 4.85 X_1$<br>$+ 10.08 X_2 + 2.59 X_1 X_2$<br>$- 1.38 X_1^2 - 14.01 X_2^2$ | | | | $Y(\text{Lipid content})$<br>$= 14.47 + 4.30 X_1 - 3.12 X_2$<br>$- 0.28 X_1 X_2 - 1.50 X_1^2 - 4.03 X_2^2$ | | | |
| Source | Estimate | p-values |  | Source | Estimate | p-values |  |
| Model | 39.26 | < e-15 | *** | Model | 14.47 | < e-15 | *** |
| X<sub>1</sub> | 4.85 | < e-08 | *** | X<sub>1</sub> | 4.30 | < e-10 | *** |
| X<sub>2</sub> | 10.08 | < e-12 | *** | X<sub>2</sub> | -3.12 | < e-07 | *** |
| X<sub>1</sub>:X<sub>2</sub> | 2.59 | < e-07 | *** | X<sub>1</sub>:X<sub>2</sub> | -0.28 | 0.480 |  |
| X<sub>1</sub><sup>2</sup> | -1.38 | < e-09 | *** | X<sub>1</sub><sup>2</sup> | -1.50 | 0.011 | * |
| X<sub>2</sub><sup>2</sup> | -14.01 | < e-08 | *** | X<sub>2</sub><sup>2</sup> | -4.03 | < e-05 | *** |
| Dependent variable: Biomass (g/L) |  |  |  |  |  |  |  |
| $Y(\text{Biomass}) = 6.12 + 0.11 X_1 + 0.36 X_2$<br>$- 0.17 X_1 X_2 - 0.02 X_1^2$<br>$- 3.91 X_2^2$ | | | | $Y(\text{Biomass}) = 3.17 + 0.71 X_1 - 0.33 X_2$<br>$- 0.13 X_1 X_2 - 0.14 X_1^2 - 0.12 X_2^2$ | | | |
| Source | Estimate | p-values |  | Source | Estimate | p-values |  |
| Model | 6.12 | < e-15 | *** | Model | 3.17 | < e-15 | *** |
| X<sub>1</sub> | 0.11 | 0.210 |  | X<sub>1</sub> | 0.71 | < e-09 | *** |
| X<sub>2</sub> | 0.36 | 0.005 | ** | X<sub>2</sub> | -0.33 | 0.005 | ** |
| X<sub>1</sub>:X<sub>2</sub> | -0.17 | 0.005 | ** | X<sub>1</sub>:X<sub>2</sub> | -0.13 | 0.238 |  |
| X<sub>1</sub><sup>2</sup> | -0.02 | 0.323 |  | X<sub>1</sub><sup>2</sup> | -0.14 | 0.344 |  |
| X<sub>2</sub><sup>2</sup> | -3.91 | < e-13 | *** | X<sub>2</sub><sup>2</sup> | -0.12 | 0.500 |  |

**Table 5.** Evaluating the significance of regression models by ANOVA.

| C. oleaginosus |  |  |  |  |  | Y. lipolytica |  |  |  |  |
| --- | --- | --- | --- | --- | --- | --- | --- | --- | --- | --- |
| Source | Lipid content % (w/w) |  |  |  |  |  |  |  |  |  |
|  | Degrees of Freedom | Sum Square | Mean Square | F-Value | p-Value | Degrees of Freedom | Sum Square | Mean Square | F-Value | p-Value |
| FO(X1, X2) | 2 | 4181.0 | 2090.50 | 356.93 | < 2.2e-16 | 2 | 562.80 | 281.40 | 141.65 | 6.459e-13 |
| TWI (X1, X2) | 1 | 497.2 | 497.20 | 84.89 | 2.374e-09 | 1 | 1.15 | 1.15 | 0.58 | 0.4561 |
| PQ ( X1, X2) | 2 | 1022.5 | 511.27 | 87.29 | 9.709e-12 | 2 | 128.23 | 64.11 | 32.27 | 3.936e-07 |
| Residual | 24 | 140.6 | 5.86 |  |  | 21 | 41.72 | 1.99 |  |  |
| Lack of fit | 4 | 82.5 | 20.63 | 7.11 | 0.000992 | 2 | 29.90 | 14.95 | 24.04 | 6.237e-06 |
| Pure error | 20 | 58.1 | 2.90 |  |  | 19 | 11.81 | 0.62 |  |  |
| Dependent variable= Lipid content % (w/w); R<sup>2</sup>= 0.9759; Adj R<sup>2</sup>= 0.9709, p-value= < e-15 |  |  |  |  |  | Dependent variable= Lipid content % (w/w); R<sup>2</sup>= 0.9432; Adj R<sup>2</sup>= 0.9296, p-value= <e-11 |  |  |  |  |
| Source | Biomass (g/L) |  |  |  |  |  |  |  |  |  |
|  | Degrees of Freedom | Sum Square | Mean Square | F-Value | p-Value | Degrees of Freedom | Sum Square | Mean Square | F-Value | p-Value |
| FO(X1, X2) | 2 | 23.91 | 11.97 | 69.94 | 1.128e-10 |  | 14.03 | 7.01 | 49.52 | 1.122e-08 |
| TWI (X1, X2) | 1 | 0.31 | 0.31 | 1.80 | 0.1919 | 1 | 0.24 | 0.24 | 1.70 | 0.2063 |
| PQ ( X1, X2) | 2 | 42.61 | 21.30 | 122.84 | 2.467e-13 | 2 | 0.16 | 0.08 | 0.55 | 0.5834 |
| Residual | 24 | 4.16 | 0.17 |  |  | 21 | 2.97 | 0.14 |  |  |
| Lack of fit | 4 | 3.43 | 0.86 | 23.58 | 2.483e-07 | 2 | 2.37 | 0.19 | 37.48 | 2.539e-07 |
| Pure error | 20 | 0.73 | 0.04 |  |  | 19 | 0.60 | 0.03 |  |  |
| Dependent variable= Biomass (g dcw/L); R<sup>2</sup>= 0.9414; Adj R<sup>2</sup>= 0.9292, p-value= <e-13 |  |  |  |  |  | Dependent variable= Biomass (g dcw/L); R<sup>2</sup>= 0.8291; Adj R<sup>2</sup>= 0.7884, p-value= <e-06 |  |  |  |  |
| FO, TWI, and PQ refer to the linear function, two-way interactions, and quadratic terms in the model formula of RSM respectively. |  |  |  |  |  |  |  |  |  |  |

ANOVA was performed to assess the significance and adequacy of response surface quadratic models. The quality of the model fit can be evaluated by the coefficient of determination (R^2^), which provides a measure of how much variability in the observed response values can be explained by the experimental factors and their interactions. The R^2^ value is always between 0 and 1 and the closer the R^2^ value is to 1, the stronger the model is and the better it predicts the response. For the developed models of *C. oleaginosus* and *Y. lipolytica*, the determination confidence coefficients (R^2^) are 97.58 % and 94.32 % for lipid content, 94.14 % and 82.91 % for biomass. These R^2^ values represented that sample variation of the regression models described the experimental data accurately. The model is regarded as significant if the p-value is lower than 0.05. In other words, ‘Model F-value’ could occur because of noise with only a 5% chance [33]. Therefore p-value of the models, for *C. oleaginosus* P_lipid accumulation_= <e-15, Pbiomass= < e-13, and for *Y. lipolytica* P_lipid accumulation_= < e-11, and P_biomass_= < e-6 suggested the coefficients are significant. The Lack of Fit P-values showed that the Lack of Fit F-value could occur due to noise with the possibility of almost 0 % for the second model for *C. oleaginosus* and 0 % for all models for *Y. lipolytica*, and 0.0992 % for the first model of *C. oleaginosus*.

### Response Surface Analysis

The factors, C/N ratio of the growth medium, and temperature were selected to be optimized for *C. oleaginosus* and *Y. lipolytica*. Three-dimensional surface responses were plotted to illustrate the relationships between the responses and variables for *C. oleaginosus* (Figure 1) and *Y. lipolytica* (Figure 2). When the temperature was around 30 °C and the C/N ratio was between C/N 75 to C/N 220, the lipid accumulation of *C. oleaginosus* was improved (Figure 1-a). On the other hand, the biomass density increased within the range of C/N 60 to C/N 240 and from 25 °C to 30 °C (Figure 1-b). Additionally, lipid accumulation of *Y. lipolytica* was enhanced starting from C/N 100 and between 18 °C to 25 °C (Figure 2-a).

**Figure 1.**
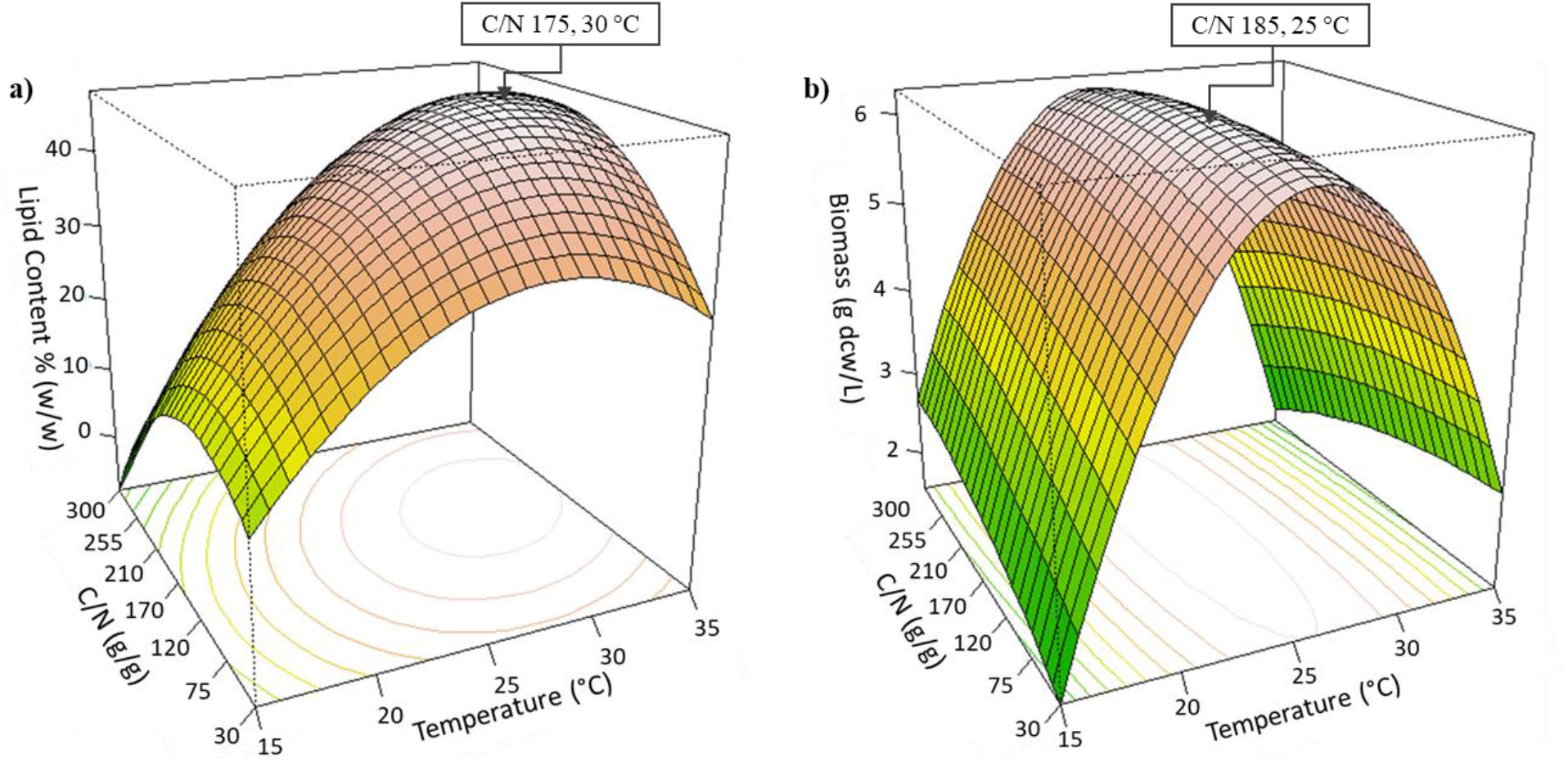
3D Response surface plot of the combined effects of C/N ratio and temperature levels on a) lipid content (g lipid weight/g yeast dry cell weight), and b) growth (g yeast dry cell weight /L) of *C. oleaginosus*. Determined optima are highlighted in the respective plots.

**Figure 2.**
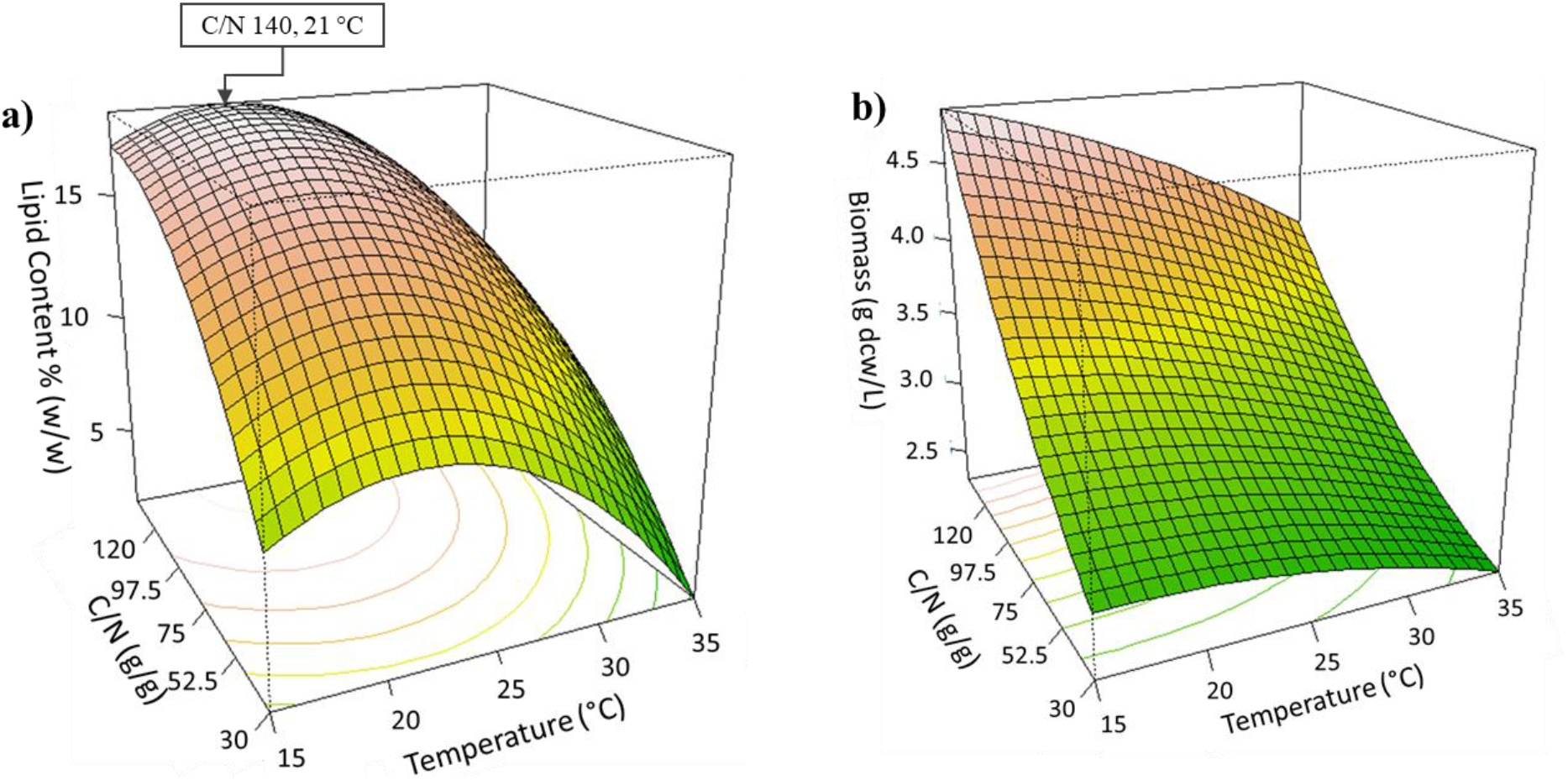
3D Response surface plot of the combined effects of C/N ratio and temperature levels on a) lipid content (g lipid weight/g yeast dry cell weight), and b) growth (g yeast dry cell weight /L) of *Y. lipolytica*. Determined optima are highlighted in the respective plots.

The optimal values for the investigated dependence factors were predicted from these 3D response surface plots (Figure 1 and Figure 2). Accordingly, the maximum predicted responses for *C. oleaginosus* were 47.67 % lipid accumulation and 6.26 g yeast dry cell weight/L. While the optimum C/N ratio and temperature were approximately C/N 175 and 30 °C for predicted lipid accumulation, C/N 185 and 25 °C were suggested to achieve the predicted maximum biomass density. The suggested optimum values for lipid accumulation resulted in 51.50 % ± 2.84 lipid accumulation, and 5.29 ± 0.08 g dry yeast cells/L which confirmed the predictions of developed regression models (Table 6). Moreover, suggestions of RSM improved lipid accumulation by 9 % in *C. oleaginosus*. Regression models of *Y. lipolytica* predicted the optimum conditions as C/N 140 and 21 °C for maximum lipid production, which predicted 18.33 % lipid accumulation, and 4.72 g yeast dry cell weight/L (Table 6). These predicted optimum conditions provided 19.03 % ± 0.37 lipid accumulation, and 5.07 ± 0.16 g dry yeast cells/L for *Y. lipolytica*. Additionally, model predictions were validated via experiments at C/N 45 and 30 °C for *C. oleaginosus* and C/N 180 at 21 °C for *Y. lipolytica* (Table 6).

**Table 6.** Validation of RSM models by suggested optimum conditions and additional experiments.

|  |  | *C. oleaginosus* |  |  |  |
| --- | --- | --- | --- | --- | --- |
|  |  | Predicted Values |  | Measured/Calculated Values |  |
| C/N (g/g) | Temperature (°C) | Lipid Accumulation (% g/g) | Biomass (g/L) | Lipid Accumulation (% g/g) | Biomass (g/L) |
| 45 | 30 | 36.08 | 5.30 | 38.48 ± 1.61 | 5.28 ± 0.08 |
| 175 | 30 | 47.67 | 4.97 | 51.17 ± 0.66 | 5.35 ± 0.28 |
|  |  | *Y. lipolytica* |  |  |  |
|  |  | Predicted Values |  | Measured/Calculated Values |  |
| C/N (g/g) | Temperature (°C) | Lipid Accumulation (% g/g) | Biomass (g/L) | Lipid Accumulation (% g/g) | Biomass (g/L) |
| 140 | 21 | 18.33 | 4.72 | 19.03 ± 0.37 | 5.07 ± 0.16 |
| 180 | 21 | 17.20 | 5.81 | 17.72 ± 0.16 | 4.96 ± 0.16 |

### Analysis of the fatty acid profile

Principal Component Analysis (PCA) was conducted to clarify the variation in the fatty acid profile at different C/N ratios and temperatures. Produced the fatty acid compositions of *C. oleaginosus* and *Y. lipolytica* are represented in Table 7 and Table 8. The PCA showing the variation in the fatty acid profile is shown in Figure 3. As seen in Figure 3-a for *C. oleaginosus* the variance explained in PC1 and PC2 were 43 % and 22 % respectively and for *Y. lipolytica* (Figure 3-b) was 56 % and 20 %. As observed from Figure 3-a, the highest temperature (35 °C) and the combination of C/N 30 with the other tested temperatures (15 °C, 30 °C) caused a higher content of saturated and longer chain fatty acids (C20:0, C22:0, C24:0). *C. oleaginosus* produced higher content of unsaturated fatty acids (C18:1, C18:2, C18:3) at the lowest temperature, 15 °C. On the other hand, we observed the same effect on the fatty acid profile of *Y. lipolytica* in terms of saturation level and chain length by changing the temperature. *Y. lipolytica* produced higher content of C14:0, C16:0, C18:0, C20:0, and C24:0 when it was incubated at 35 °C, and C14:1, C16:1, C18:1, and C20:1 at 15 °C.

**Figure 3.**
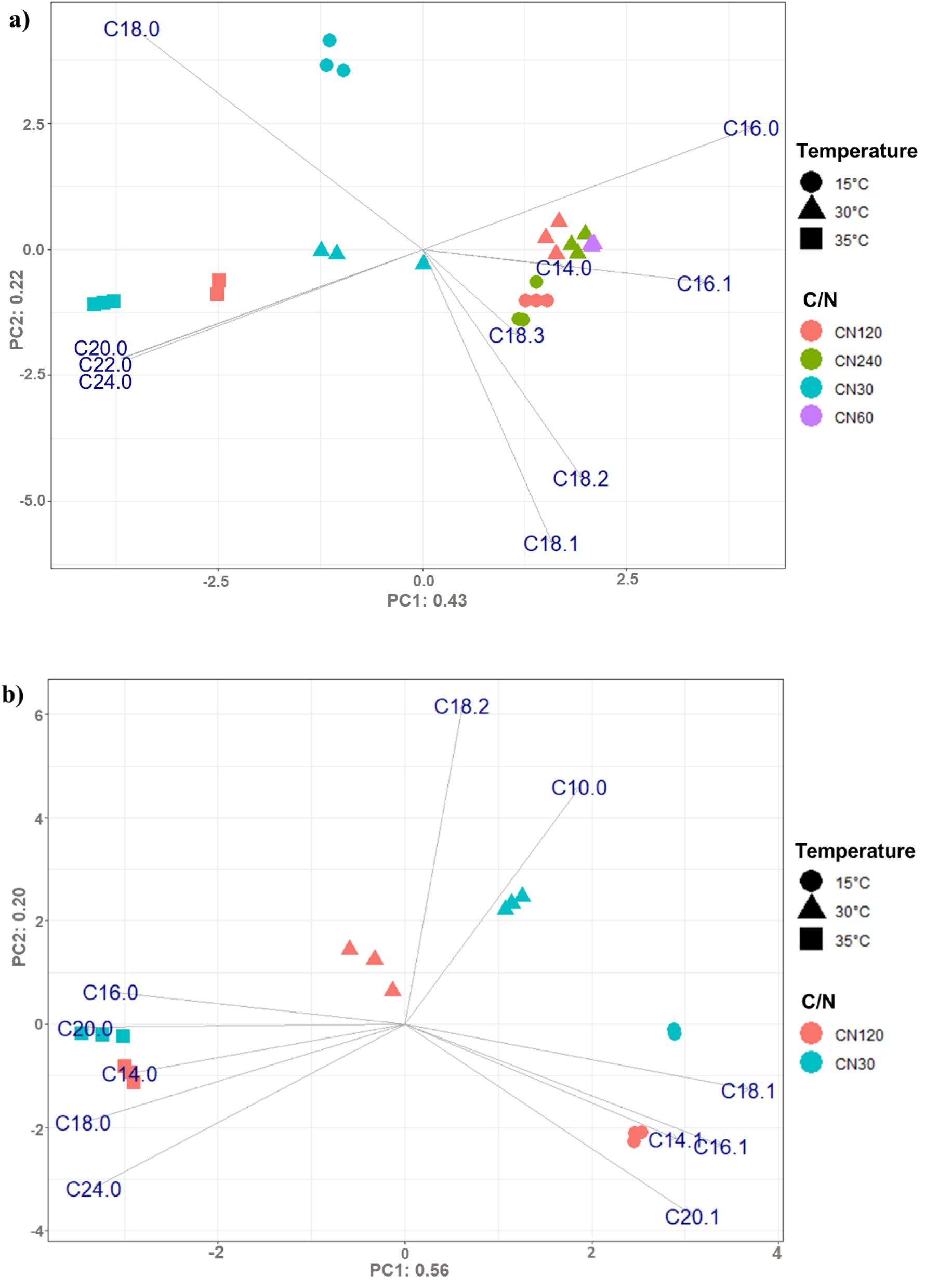
PCA on the fatty acid profile at different C/N ratios in the growth medium and temperatures for **a**)*C. oleaginous* and **b)***Y. lipolytica*. Variance explained by each component is given in the labels.

**Table 7.** Fatty acid profile of *C. oleaginosus* at various C/N ratios and temperatures.

|  |  | Fatty Acid Profile (%) |  |  |  |  |  |  |  |  |  |  |  |  |  |  |
| --- | --- | --- | --- | --- | --- | --- | --- | --- | --- | --- | --- | --- | --- | --- | --- | --- |
| C/N (g/g) | Temperature (°C) | C14:0 | C16:0 | C16:1 | C18:0 | C18:1 | C18:2 | C18:3 | C20:0 | C22:0 | C24:0 | Saturated FAs | MUFAs | PUFAs | Long chain FAs | Very long chain FAs |
| 30 | 15 | - | 24.62 ± 1.47 | - | 43.92 ± 0.53 | 26.58 ± 1.09 | 4.88 ± 1.96 | - | - | - | - | 68.54 ± 1.47 | 26.58 ± 1.09 | 4.88 ± 1.96 | 100 ± 0.0 | - |
|  | 30 | - | 15.44 ± 3.06 | - | 21.09 ± 4.22 | 46.70 ± 1.41 | 14.27 ± 1.93 | - | - | - | 2.50 ± 0.54 | 39.03 ± 1.05 | 46.7 ± 1.41 | 14.27 ± 1.93 | 97.5 ± 0.54 | 2.5 ± 0.54 |
|  | 35 | - | 9.33 ± 0.58 | - | 28.14 ± 0.80 | 42.52 ± 0.86 | 12.50 ± 0.81 | - | 1.90 ± 0.17 | 1.27 ± 0.06 | 4.34 ± 0.06 | 44.98 ± 0.05 | 42.52 ± 0.86 | 12.50 ± 0.81 | 94.39 ± 0.0 | 5.61 ± 0.0 |
| 60 | 30 | - | 21.00 ± 1.67 | 1.09 ± 0.07 | 18.96 ± 2.74 | 51.06 ± 0.87 | 8.25 ± 0.91 | - | - | - | - | 39.96 ± 2.26 | 52.15 ± 1.36 | 8.25 ± 0.91 | 100 ± 0.0 | - |
| 120 | 15 | - | 18.32 ± 0.54 | 1.79 ± 0.43 | 10.74 ± 0.63 | 56.16 ± 1.64 | 10.74 ± 0.62 | 2.26 ± 0.24 | - | - | - | 29.05 ± 0.24 | 57.95 ± 1.21 | 13.00 ± 0.97 | 100 ± 0.0 | - |
|  | 30 | 0.39 ± 0.03 | 30.33 ± 2.14 | 2.71 ± 0.23 | 7.39 ± 0.49 | 46.50 ± 1.98 | 11.93 ± 0.50 | - | 0.44 ± 0.01 | - | 0.30 ± 0.02 | 38.85 ± 2.57 | 49.22 ± 2.14 | 11.93 ± 0.50 | 99.7 ± 0.02 | 0.30 ± 0.02 |
|  | 35 | 0.40 ± 0.01 | 14.61 ± 0.11 | 0.71 ± 0.01 | 22.09 ± 0.22 | 47.85 ± 0.04 | 8.54 ± 0.18 | - | 1.75 ± 0.06 | 1.13 ± 0.03 | 3.09 ± 0.00 | 43.34 ± 0.43 | 48.55 ± 0.03 | 8.54 ± 0.18 | 95.95 ± 0.08 | 4.22 ± 0.04 |
| 240 | 15 | - | 24.89 ± 2.20 | - | - | 59.08 ± 2.55 | 16.01 ± 0.82 | - | - | - | - | 24.89 ± 2.20 | 59.08 ± 2.55 | 16.01 ± 0.82 | 100 ± 0.0 | - |
|  | 30 | 0.38 ± 0.03 | 29.30 ± 1.46 | 2.88 ± 0.10 | 6.82 ± 0.19 | 47.23 ± 1.49 | 13.10 ± 0.33 | - | - | - | 0.29 ± 0.03 | 36.79 ± 1.62 | 50.11 ± 1.51 | 13.10 ± 0.33 | 100 ± 0.0 | 0.29 ± 0.03 |
| MUFAs: Monounsaturated fatty acids, PUFAs: Polyunsaturated fatty acids |  |  |  |  |  |  |  |  |  |  |  |  |  |  |  |  |

**Table 8.** Fatty acid profile of *Y. lipolytica* at various C/N ratios and temperatures.

|  |  | Fatty Acid Profile (%) |  |  |  |  |  |  |  |  |  |  |  |  |  |  |  |
| --- | --- | --- | --- | --- | --- | --- | --- | --- | --- | --- | --- | --- | --- | --- | --- | --- | --- |
| C/N (g/g) | Temperature (°C) | C10:0 | C14:0 | C14:1 | C16:0 | C16:1 | C18:0 | C18:1 | C18:2 | C20:0 | C20:1 | C24:0 | Saturated FAs | MUFAs | PUFAs | Long chain FAs | Very long chain FAs |
| 30 | 15 | 1.27 ± 0.15 | - | 0.68 ± 0.01 | 8.56 ± 0.34 | 13.41 ± 0.37 | 6.27 ± 0.70 | 46.54 ± 1.17 | 21.41 ± 0.01 | - | 1.84 ± 0.32 | - | 16.11 ± 1.20 | 62.48 ± 1.22 | 21.41 ± 0.01 | 98.72 ± 0.16 | - |
|  | 30 | 2.53 ± 0.31 | - | - | 10.59 ± 0.35 | 11.47 ± 0.68 | 5.57 ± 0.38 | 43.97 ± 0.88 | 24.67 ± 0.44 | 1.18 ± 0.10 | - | - | 19.88 ± 0.27 | 55.44 ± 0.21 | 24.67 ± 0.44 | 97.47 ± 0.31 | - |
|  | 35 | - | 1.02 ± 0.31 | - | 18.77 ± 0.27 | 6.47 ± 0.18 | 14.94 ± 1.37 | 33.83 ± 0.13 | 19.14 ± 0.71 | 3.60 ± 0.27 | - | 2.22 ± 0.04 | 40.56 ± 1.02 | 40.30 ± 0.31 | 19.14 ± 0.71 | 97.77 ± 0.04 | 2.22 ± 0.04 |
| 120 | 15 | - | - | 0.30 ± 0.00 | 10.26 ± 0.10 | 19.98 ± 0.66 | 5.38 ± 0.25 | 50.21 ± 0.43 | 11.38 ± 0.20 | - | 1.81 ± 0.03 | 0.68 ± 0.01 | 16.32 ± 0.40 | 72.29 ± 0.52 | 11.38 ± 0.20 | 99.32 ± 0.10 | 0.68 ± 0.01 |
|  | 30 | - | - | - | 23.31 ± 0.58 | 11.40 ± 1.37 | 7.81 ± 0.66 | 35.73 ± 1.06 | 21.74 ± 1.39 | - | - | - | 31.12 ± 0.08 | 47.14 ± 1.30 | 21.75 ± 1.38 | 100 ± 0.00 | - |
|  | 35 | - | 0.29 ± 0.01 | - | 20.97 ± 0.05 | 6.79 ± 0.24 | 22.14 ± 1.06 | 32.75 ± 1.23 | 12.72 ± 0.58 | 2.23 ± 0.14 | - | 2.11 ± 0.13 | 47.73 ± 0.87 | 39.54 ± 1.42 | 12.72 ± 0.58 | 97.88 ± 0.13 | 2.11 ± 0.13 |
| MUFAs: Monounsaturated fatty acids, PUFAs: Polyunsaturated fatty acids |  |  |  |  |  |  |  |  |  |  |  |  |  |  |  |  |  |

## Discussion

The lipid accumulation and lipid productivity in oleaginous yeasts are strongly dependent on the composition of the cultivation medium and the operational conditions. Therefore, this study sought to assess the importance of such factors in lipid production. To this end, we performed an RSM analysis and represented the optimum C/N ratio and temperature for the maximization of lipid content, and biomass density. The optimum C/N ratio and temperature are C/N 175 at 30°C for *C. oleaginosus* and C/N 140 at 21°C for *Y. lipolytica*. Moreover, we demonstrated that a C/N ratio and temperature cause variations in the fatty acid composition of oleaginous yeasts.

In the experiments performed for RSM, we used glycerol as a carbon source as this is efficiently utilized by *C. oleaginosus* and *Y. lipolytica* [34,35] and urea as a nitrogen source as it provides higher biomass yields compared to the ammonium salts [26]. RSM is one of the most preferred methods to optimize operational conditions and medium composition in biotechnology [36,37]. This method facilitates obtaining more information with a fewer number of experiments by changing multiple factors at a time because RSM reflects on the complex nonlinear relationships between independent variables and measured responses of the system. In contrast, the OFAT approach allows changing only one of the factors for each of the experiments. Whereas around 15 experiments are required to follow the OFAT approach with the same factors and same levels, the number of experiments decreased by approximately 40 % via the DoE approach. The statistics tables of developed regression models in this study represented that both the C/N ratio and the temperature have a significant effect on the lipid content of cells, and biomass density of *Y. lipolytica*. These results are similar to those reported by Canonico et al [17]. Although they utilized crude glycerol as a cultivation medium, the behavior of biomass and lipid content against changing temperature and C/N ratio is comparable with the results in this study. On the contrary, in this study, the growth of *C. oleaginosus* was significantly affected by only temperature and the combined effect of temperature and C/N ratio. These findings are consistent with the report of Cui et al. even though they tested a narrower temperature range between 27 °C to 33 °C, whereas in our design, it was extended from 15 °C to 35 °C [27]. Although the maximum lipid accumulation of *Y. lipolytica* is much less than *C. oleaginosus*, it is corresponding to the amounts reported for the wild-type strain [38,39]. Gao et al. reported lipid accumulation of *Y. lipolytica* CICC 31596 up to 30 % (g/g) when it grew on volatile fatty acids [40]. On the other hand, *Y. lipolytica* ACA-DC 50109 produced 20 % (g/g) lipids on a glycerol-based cultivation medium [34]. These findings show lipid accumulation of *Y. lipolytica* is strain-dependent. On the other hand, optimum temperature, 21 °C, for lipid accumulation and growth for *Y. lipolytica* identified in this study was surprisingly lower than previous reports. Our findings provided an optimum C/N ratio and temperature as these predicted values provided higher lipid contents, and biomass for both oleaginous yeasts. These optimum values identified via RSM in this study will potentially contribute to solving other optimization problems such as revealing the effect of other factors, and finding optimum conditions for other strains and engineered strains. In addition to optimization of cultivation conditions, lipid accumulation ability of *Y. lipolytica* and *C. oleaginosus* can be further improved by strain engineering as suggested in other reports [41].

In addition to the lipid content and biomass, the C/N ratio and temperature also affected the fatty acids composition of *C. oleaginosus* and *Y. lipolytica*. In previous studies, Moon et al. reported that 15 °C as a growth temperature shifted the fatty acid profile of *C. oleaginosus* to more unsaturated fatty acids [42]. Moreover, Hackenschmidt et al. claimed that there was only a slight variation in the fatty acid profile of *Y. lipolytica* from 25 °C to 35 °C [43]. However, systematic evaluation of the low and high levels of temperature and C/N ratio have not been performed to our knowledge. In this study, PCA allowed us to evaluate the variation in produced fatty acids composition by reducing the noise and creating uncorrelated components from the analyzed data. When the growth temperature of oleaginous yeasts was increased from optimum to 35 °C and the C/N ratio decreased to C/N 30, the saturation level and the chain length of fatty acids were increased. On the other hand, fatty acids produced at 15 °C were slightly more unsaturated (C14:1, C16:1, C18:1, and C20:1 for *Y. lipolytica*, C18:1, C18:2, and C18:3 for *C. oleaginosus*) than they were at the optimum temperature. This variety can be explained by the adaptation of an organism to maintain lipid fluidity at different temperatures [42]. Because the saturation level and the chain length directly influence the melting point of fatty acids. The melting points are much lower for an unsaturated and shorter chain length of fatty acids whereas it is higher for saturated and longer chain fatty acids [44]. While low incubation temperatures increased unsaturation levels on both oleaginous yeasts, it decreased the growth and lipid productivity of *C. oleaginosus*. Unexpectedly, low incubation temperatures positively affected lipid accumulation and biomass production in *Y. lipolytica*. This situation demonstrated that *Y. lipolytica* is promising to produce fatty acids, especially with higher content of unsaturated fatty acids as growth and lipid accumulation were enhanced by lower temperature. Tezaki et al. related functions of some genes in *Y. lipolytica* with the adaptation ability of this organism at low temperatures [45]. Therefore, elucidation of the adaptation mechanism of oleaginous yeasts to low temperatures or too high temperatures could contribute to achieving higher productivity and expanding the application potential of microbial oils.

## Conclusion

In this study, we sought to determine the major operational factors affecting physiological fitness toward fatty acid production by oleaginous yeasts. We aimed to enhance lipid accumulation as well as to enable *C. oleaginosus* and *Y. lipolytica* to attain higher content of particular fatty acids by changing operating conditions such as the available C/N ratio and temperature. We applied a thorough DoE method (RSM) and developed second-order polynomial equations to identify the optimum C/N ratios and temperatures for both oleaginous yeasts. The predictions of RSM improved the lipid accumulation by approximately 71 % for *C. oleaginosus* and about 66 % for *Y. lipolytica* compared to the average lipid accumulation in the tested conditions. While the lipid accumulation was significantly affected by the C/N ratio and temperature, the growth of *C. oleaginosus* was mainly affected by temperature. Additionally, changing the C/N ratio in the cultivation medium and temperature resulted in variations in fatty acid profile, which we observed switches from saturated to unsaturated fatty acids, unsaturated to saturated fatty acids, and shorter chain to longer chain fatty acids. Altogether, these findings helped strengthen the basis to deploy these oleaginous yeasts as platforms for tailored fatty acid production and thereby contribute to the development of processes substituting palm oil that are more sustainable.

## Supplementary Material

**Figure S1.**
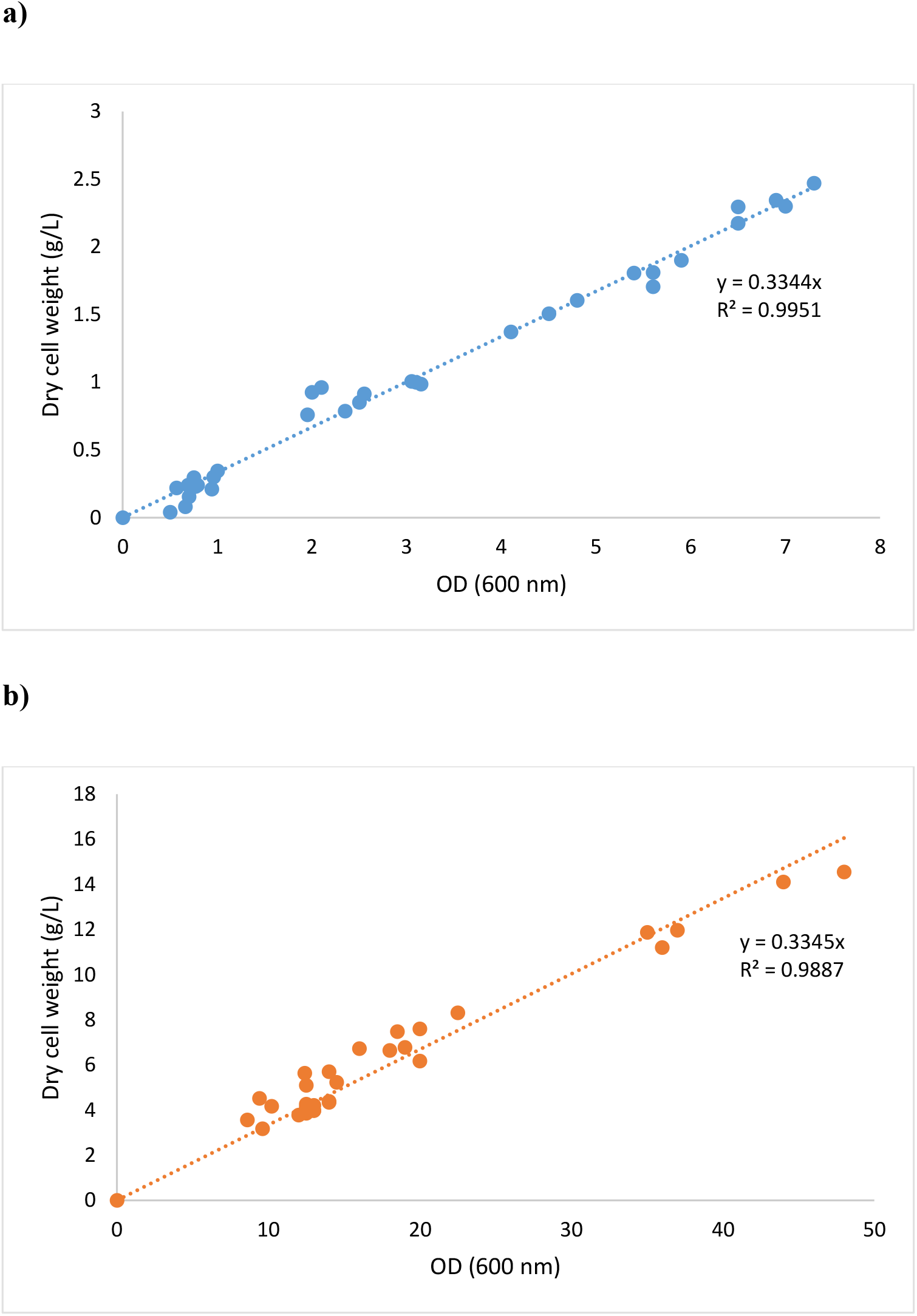
OD versus dry cell weight curve for a) *C. oleaginosus* and b) *Y. lipolytica*.

## Author’s contributions

All authors conceived and designed the study. ZEDÖ and MSD performed the data analysis. ZEDÖ drafted the manuscript and performed the experiments. VAPMdS, JH, and MSD acquired project funding, conceived and supervised the research. All authors reviewed and edited the study. All authors read and approved the final manuscript.

## Acknowledgments

We thank Sara Moreno Paz for her valuable contribution to design of experiment and for providing the initial code for RSM. We thank Janine Verbokkem for her valuable contribution to the analysis of fatty acid composition.

## Author details

^1^Bioprocess Engineering, Wageningen University & Research, 6708 PB, Wageningen, the Netherlands. ^2^Laboratory of Systems and Synthetic Biology, Wageningen University & Research, 6708 WE, Wageningen, the Netherlands. ^2^LifeGlimmer GmbH, Berlin, 12163, Germany. ^4^Wageningen Food & Biobased Research, Wageningen University & Research, 6708 WE, Wageningen, The Netherlands.

